# A novel histone H4 variant regulates rDNA transcription in breast cancer

**DOI:** 10.1101/325811

**Authors:** Mengping Long, Xulun Sun, Wenjin Shi, Yanru An, Tsz Chui Sophia Leung, Dongbo Ding, Manjinder S. Cheema, Nicol MacPherson, Chris Nelson, Juan Ausio, Yan Yan, Toyotaka Ishibashi

## Abstract

Histone variants, present in various cell types and tissues, are known to exhibit different functions. For example, histone H3.3 and H2A.Z are both involved in gene expression regulation, whereas H2A.X is a specific variant that responds to DNA double-strand breaks. In this study, we characterized H4G, a novel hominidae-specific histone H4 variant. H4G expression was found in a variety of cell lines and was particularly overexpressed in the tissues of breast cancer patients. H4G was found to localize primarily to the nucleoli of the cell nucleus. This localization was controlled by the interaction of the alpha helix 3 of the histone fold motif with the histone chaperone, nucleophosphomin 1. In addition, we found that H4G nucleolar localization increased rRNA levels, protein synthesis rates, and cell cycle progression. Furthermore, micrococcal nuclease digestion of H4G-containing nucleosomes reconstituted *in vitro* indicated that H4G destabilizes the nucleosome, which may serve to alter nucleolar chromatin in a way that enhances rDNA transcription in breast cancer tissues.

## Introduction

Eukaryote genomic DNA is tightly packaged into a nucleoprotein structure called chromatin, which consists of discrete nucleosome subunits. The nucleosome core particle is further composed of 147 DNA base pairs (bp) wrapped around a histone core octamer. Each histone octamer comprises two copies of the core histones H2A, H2B, H3, and H4. In addition to the core histones, histone H1 is a linker histone that binds to linker DNA, connecting adjacent nucleosomes in close proximity to the canonical histone octamer region, helping to further compact nucleosomal DNA in the chromatin fiber [1]. Epigenetic markers affecting chromatin structure include histone variants and histone post-translational modifications (PTMs), which regulate important biological functions, including DNA replication and repair, and transcription [2].

Epigenetic alterations, such as changes in histone PTMs and DNA methylation, are integral to regulating gene expression patterns and maintaining genome stability [3–5]. Changes in the expression patterns of histone variants have been reported to be related to the genome instability observed in cancer cells [6]. For example, H2A.X phosphorylation is essential for the repair of DNA double-strand breaks, and H2A.X-knockout mice show immunodeficiency, radiosensitivity, and a high susceptibility to cancer [7–9]. MacroH2A operates as a transcription repressor involved in X-chromosome inactivation [10] and the level of MacroH2A expression is decreased in both breast and colon cancers [11]. MacroH2A is also believed to be a senescence marker, with the loss of macroH2A contributing to tumor progression through a bypass of senescence [11, 12]. H2A.Z expression levels are increased in many cancer types, including colorectal, breast, and prostate cancers [11, 13]. Although increasing evidence suggests the potential link between histone variant expression and tumor progression, the mechanisms by which histone variants control cellular proliferation and growth remains poorly understood.

Many types of histone variants are present in humans, including H2A.Z, H2A.X, H2A.Bbd, macroH2A, TH2B, H2BFWT, H3.3, H3.4, H3.5, H1t, H1.X, and H1foo. Each variant has been shown to have a specific localization and expression pattern, indicating that they are functionally unique [16, 17]. Holmes et al. studied the expression of histone H4 genes [18] and found that, compared with histones H1, H2A, H2B, and H3, histone H4 is the only histone that does not have functionally distinct variants. However, in this study, we describe the function of a previously unrecognized H4 variant: H4G. The H4G protein lacks the C-terminal tail region of the canonical human histone H4 and shares only 85% identity with H4 in the remaining 98 amino acids. The gene *h4g* is located in the histone cluster 1, with many other histone genes encoded nearby, and is present only in hominids [18]. We found that *h4g* is expressed in the human breast cancer cell line MCF7 but not in the normal breast epithelial cell line MCF10A or other cell types commonly used in the lab, such as HeLa and HEK293T cells. Moreover, we found that *h4g* is expressed in breast tissues from human breast cancer patients but was not expressed in healthy breast tissue. Importantly, H4G is primarily localized to the nucleoli, and its expression positively regulates the transcription of rDNA. H4G depletion in MCF7 breast cancer cells decreased cellular proliferation rates as a result of reduced rRNA and protein synthesis. Using *in vitro* assays we demonstrated that H4G destabilizes nucleosomes. Finally, consistent with a nucleolar function for this histone variant, we identify NPM1 (nucleophosmin1), a nucleolar histone chaperone involved in ribosomal biogenesis and in cancer [19–22], as an H4G interacting protein that preferentially recognizes the α-helix 3 of the H4G histone fold domain.

## Results

### H4G expression in breast cancer cell lines and tissues

The human genome contains three histone clusters where multiple copies of the major core and linker histones as well as many histone variants are located [17]. The canonical histone genes encode proteins consisting of highly conserved amino acid sequences, and they are expressed primarily during the S phase of the cell cycle. *h4g* is encoded in histone cluster 1, which is located at the chromosome 6p22.1–22.2 region in the human genome [18]. In this study, we searched the genomes of other species, and we were only able to identify the *h4g* in hominid genomic DNA (Figure 1A). H4 is known as one of the most evolutionarily conserved histones [23, 24]. Surprisingly H4G shares only 85% amino acid identity with human canonical H4 (Figure 1A), lacking the last five amino acids of the C-terminal tail and possessing several different amino acids within the N-terminal tail and the α-helix 1, 2, and 3 domains of the histone fold (Figure 1A). In addition, compared with its canonical histone counterpart, H4G has a higher hydrophobicity, which affects amino acids in the N-terminal tail and α-helix 3 of the core domain (Figures 1A and EV1).

**Figure 1.**
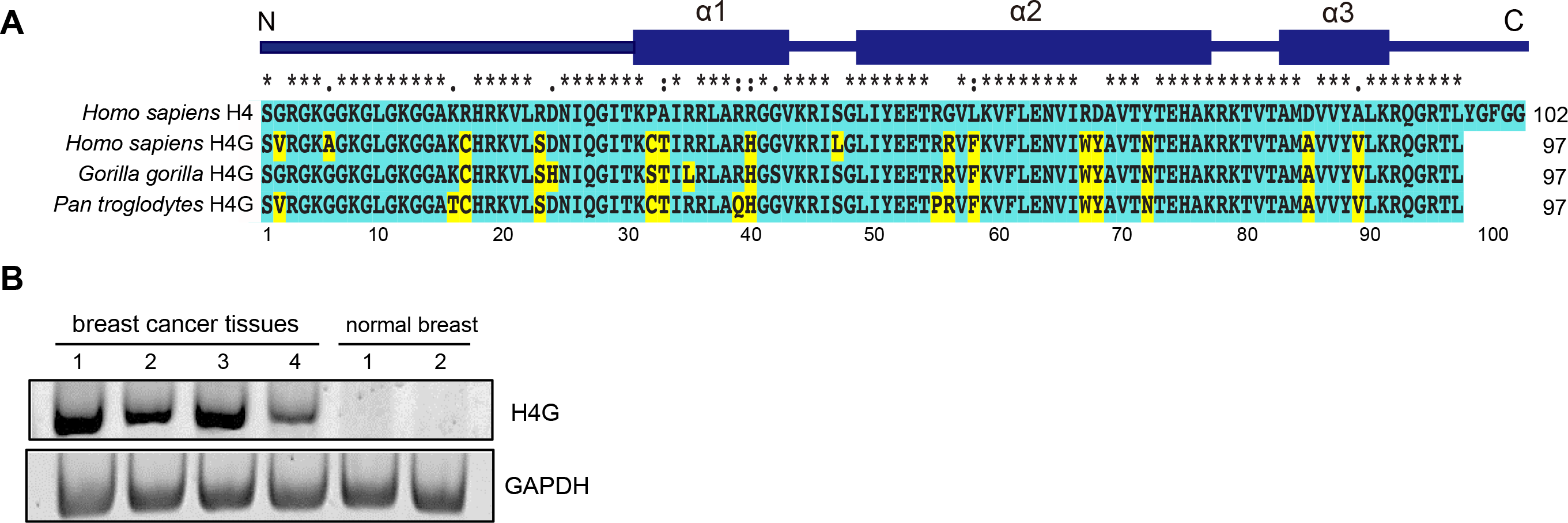
H4G expression in breast cancer patient tissues. (A) Amino acids sequence alignment between primate histone H4G and canonical H4 in primates. Unique amino acids to H4G in comparison with canonical human H4 are highlighted in yellow. (B) The H4G expression in human breast cancer patient breast tissues and normal breast tissues. Detailed sample origins are as follows: samples 1-4 were human breast adenocarcinoma RNA purchased from ORIGENE (catalog number CR560317, 560615, 560023, 560116); sample 5 was normal breast tissue RNA (catalog number CR559173) from ORIGENE; sample 6 was normal breast RNA pool from five donors (lot number 8812195) from Biochain. The negative control for all samples failed to amplify H4G, thus the contamination of genomic DNA is excluded.

To study the function of H4G, we first checked *h4g* expression levels in several cultured cell lines. We found that *h4g* is expressed in breast cancer cell lines, including MCF7, LCC1, and LCC2 cells (Table 1). The expression of *h4g* was not detect by qPCR in the noncancerous breast epithelial cell line, MCF10A; the cervical cancer cell line, HeLa; the lung cancer cell lines, H1299 and PC9; or the embryonic kidney cell line, HEK293T (Table 1). Although *h4g* expression levels in breast cancer cell lines were generally low and the low expression in cultured cells agrees with the expression data from database (The Human Protein Atlas), the *h4g* expression level was higher in the tamoxifen-resistant cell line LCC2 than in the tamoxifen-sensitive cell line LCC1, even though they are derived from the same parental cell line (MCF7)(Table 1). We also evaluated the expression of *h4g* in tissues from breast cancer patients using cDNA from those patients (ORIGENE) and compared *h4g* expression with that in normal breast and testes tissues (Zyagen and ORIGENE; Figure 1B). Interestingly, the *h4g* gene was only expressed in breast cancer patient-derived samples (4 out of 4, *N* = 4), and *h4g* expression was not detected in either normal breast (0 out of 2, *N* = 2, one of the samples was the mixture of five donors) (Figure 1B) or testes samples (data not shown).

**Table 1.**
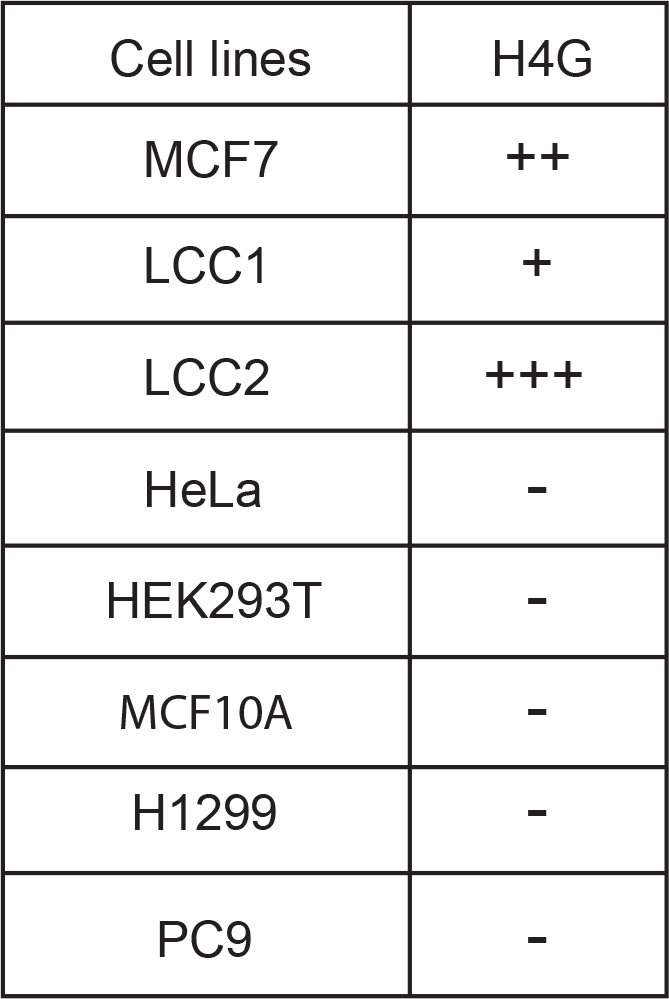
H4G expression in cultured cell

### The H4G histone fold destabilizes the nucleosome

Histone variants are known to affect nucleosomal stability; such effects underlie their regulatory role in processes such as transcription and DNA repair [25]. The C-terminal end of the H4G amino acid sequence was found to be different from that of the canonical H4, with the latter forming a β-sheet structure with H2A [26]. Furthermore, the H4G sequence was also found to be different from that of the N-terminal tail of H4, which also directly interacts with H2A [27–29]. These sequence differences indicate that H4G likely affects H4-H2A interactions and the overall stability of the nucleosome. Therefore, we analyzed the stability of reconstituted nucleosomes consisting of this variant and the 601 nucleosome positioning sequence (Figure 2A and 2B) [30]. However, the recombinant expression of H4G served to be a challenge likely due to its high hydrophobicity. As shown in Figure 2, H4G appears capable of forming a nucleosome, with the H4G-containing nucleosomes being more sensitive to the MNase treatment (Figure 2A and 2B). Hence, the different functions of H4G compared with the canonical H4 may be a direct result of their sequence differences, which differentially affect nucleosome stability.

**Figure 2.**
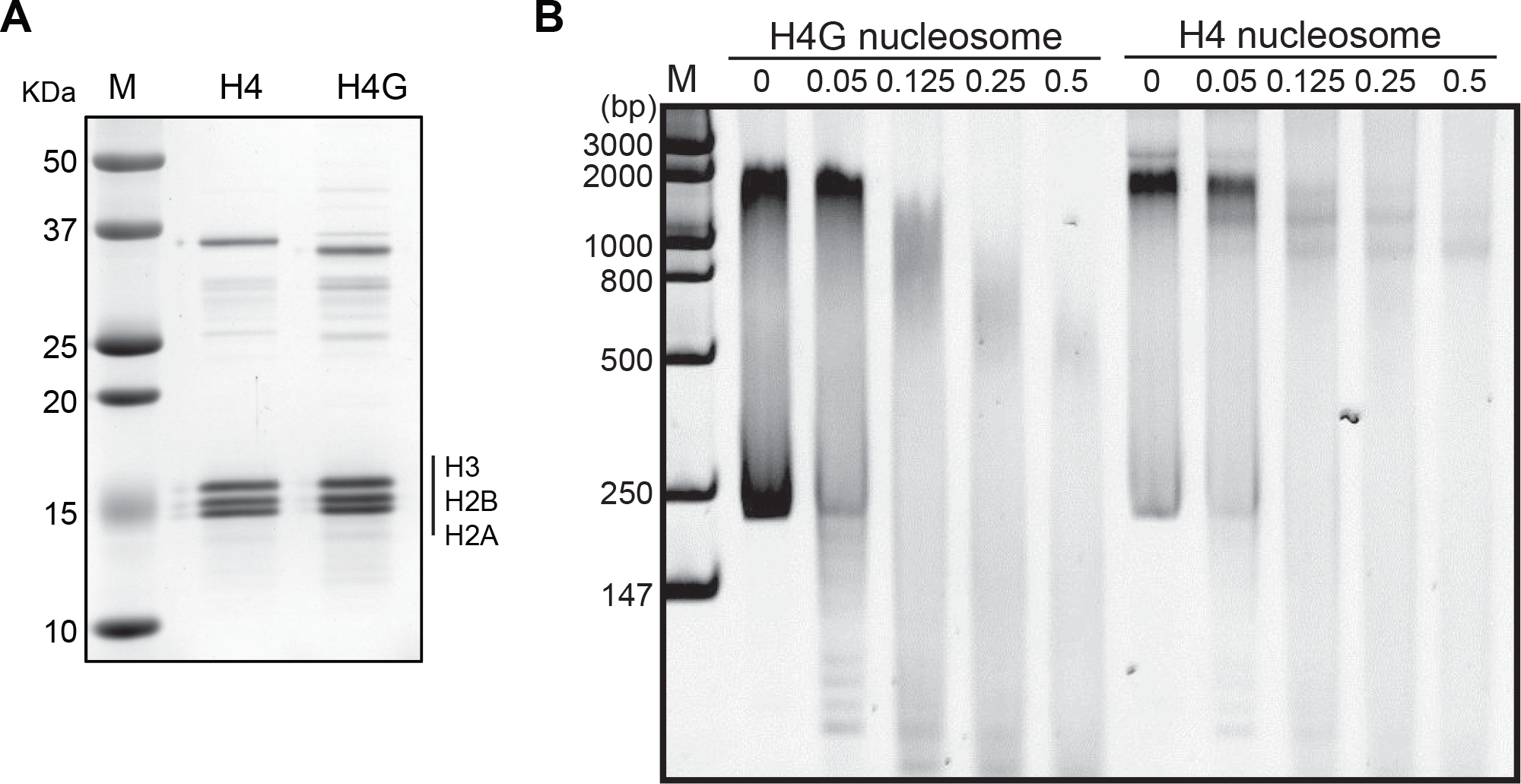
The H4G-containing nucleosome is more sensitive to MNase. (A) SDS-PAGE of the canonical histone mixture and the H4G mixture for the nucleosome loading. Asterisk (*) represents the H4-GST and H4G-GST proteins. (B) Representative polyacrylamide gel of nucleosome samples after MNase treatment.

### The α-helix 3 domain of H4G is responsible for its nucleolar localization

The different nuclear localization pattern of histone variants can be an indication of their distinct regulatory roles in different cellular processes. For instance, CENP-A localizes to the centromeric region, whereas H3.3 localizes to the promoters of active genes [25]. We found that when FLAG-tagged H4G was expressed in MCF7 cells, it localized to the nucleolus, whereas the overexpressed FLAG-tagged canonical H4 was localized to the nucleus, as expected (Figure 3A). This localization pattern was also observed in LCC1 and LCC2 cells (Figure 3B and 3C) and was not affected by the N-or C-terminal FLAG or EGFP, as was observed in other cell lines such as MCF10A, HeLa, and HEK293T (data not shown).

**Figure 3.**
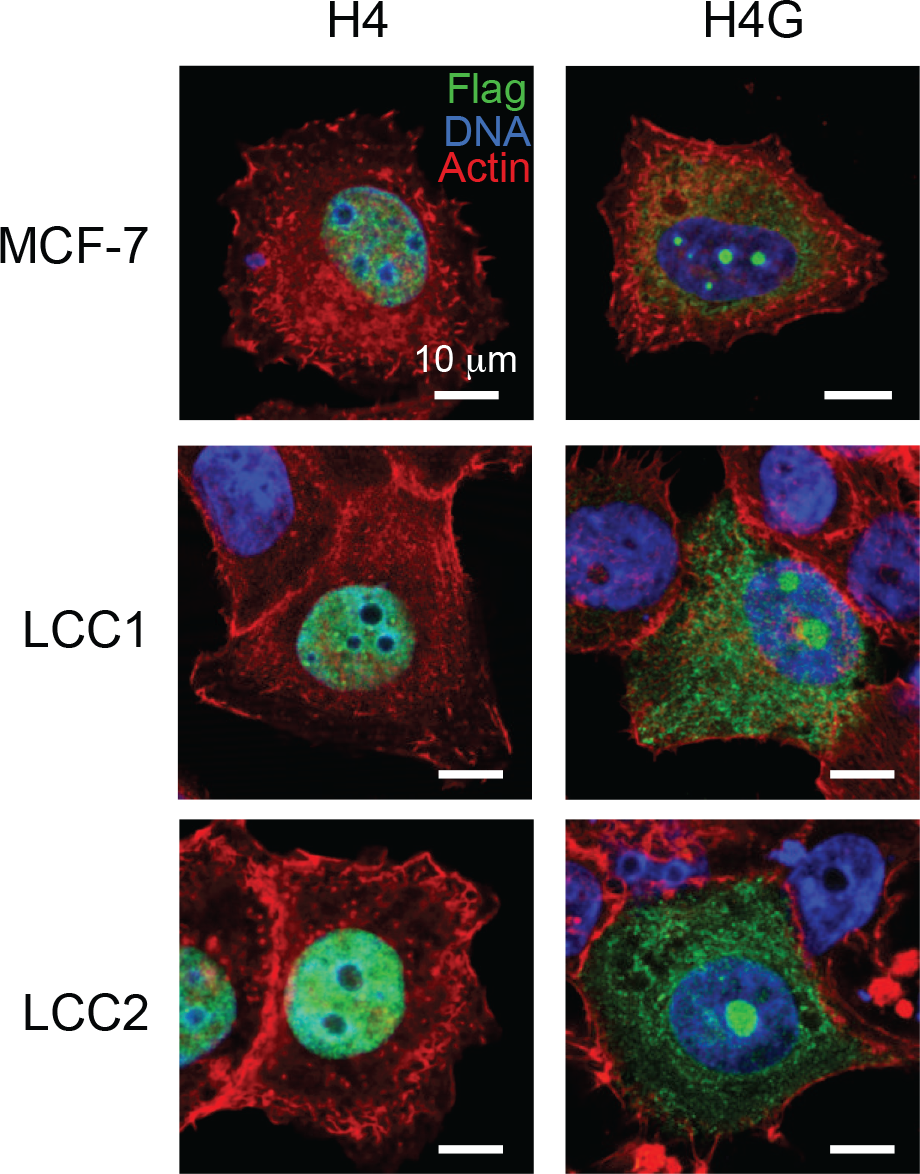
H4G localizes to the nucleolar in breast cancer cells. (A–C) The cellular localization of H4 and H4G in MCF7 (A), LCC1 (B), and LCC2 (C) cells. The scale bars represent 10 μm.

To further dissect the structural determinants of the H4G nucleolar localization, we made a series of H4G mutants, in which we swapped different H4 and H4G regions and analyzed their localization patterns (Figure 4A-4H). To study the effect of the N-terminal tail, we made constructs containing the H4 histone fold with the H4G N-terminal tail (H4sN; s stands for swapping) and a construct containing the H4G histone fold with the H4 N-terminal tail (H4GsN) (Figures 3 and 4A). The H4sN was found to still localize to the nucleus like H4, whereas the H4GsN was found to localize to the peripheral region of the nucleolus in comparison with the uniform nucleolar localization of H4G (Figures 4B and EV2). For the analysis of the C-terminal tail, we made constructs of H4sC with a deletion of the last five amino acids of the original H4 construct and H4G with both N-and C-terminal tails of H4 (H4GsN+C). The H4sC localization was similar to that of H4, whereas that of H4GsN+C localization was similar to that of H4GsN (Figure 4A, 4B, and 4D). These results indicate that the C-terminal tail of H4G does not affect on nucleolar localization. In addition, the H4 histone containing the H4G α-helix 3 domain protein (residues 85 and 89) exhibited a nucleolar localization similar to that of H4G (Figure 4G). These findings were confirmed using reverse-swapping in which the α-helix 3 of H4G was replaced with the α-helix 3 of the canonical H4, leading to the complete absence of nucleolar localization (Figure 4H).

**Figure 4.**
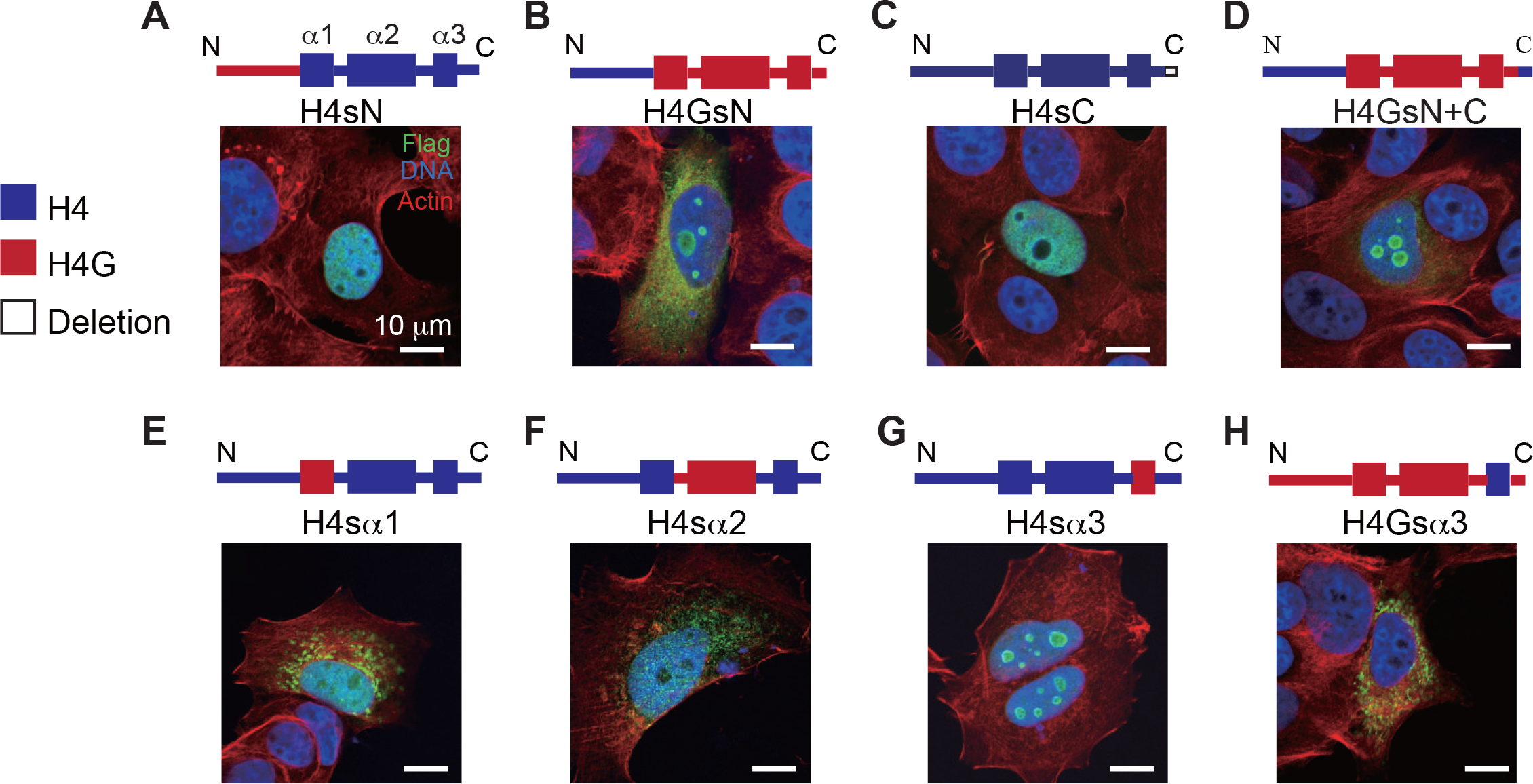
The α-helix 3 of H4G is important for the nucleolar localization. (A–H) The cellular localizations of the H4G/H4 hybrid constructs in MCF7 cells. The scale bars represent 10 μm.

The localization of H4G in the nucleolus is likely mediated by other proteins such as histone chaperones. Accordingly, we next conducted an anti-FLAG immunoprecipitation assay using HEK293T cells, which overexpress FLAG-tagged H4 or H4G, and analyzed interacting proteins by LC-MS. By comparing the interacting proteins of H4 and H4G, we found that the nucleolar histone chaperone protein, NPM1 (nucleophosmin/B23), specifically interacts with H4G but not with H4 (Table 2) [22]. A further immunoprecipitation study showed that NPM1 interacted more strongly with H3 compared with H4, which agrees with a previous report [22] (Figure EV3). However, it appears to interact more strongly with H4G, and this interaction is likely mediated by the α-helix 3 in H4G, as suggested by the disappearance of the interaction in H4Gsα3 and the presence of the interaction in H4sα3 (Figure 5). Because amino acids 85 and 89 are different in the α-helix 3 between H4 and H4G, we repeated the immunoprecipitation study to determine the role of these amino acids. As such, both positions are likely important for the interaction of H4G with NPM1, but the position 85 (D in H4 and A in H4G) appears to contribute to the interaction more than position 89 (A in H4 and V in H4G) (Figure 5). Another H3/H4 binding chaperone, Asf1, has also been documented to bind to both H3 and H4 [31, 32]. The C-terminal region of histone H4 has been shown to be necessary to function with Asf1 from H3/H4 tetramers [26]. Because H4G lacks the last five amino acids at its C-terminal end, we examined whether H4G would be able to interact with Asf1. Immunoprecipitation and mass spectrometry analyses showed that Asf1 interacts only with H4 but not with H4G (Table 2 and Figure EV4). Rbap46 and Rbap48 have also been shown to interact with the α-helix 1 of H4 [33]. In this study, we found that Rbap48 was able to interact with both H4 and H4G; however, the mass spectrometry signal of H4G was much lower than that of H4 (Table 2). Collectively, these results indicate that the α-helix 3 of H4G has an important role in its interaction with NPM1, which may be critical for its nucleolar localization.

**Table 2.** Proteins interacting with H4G and H4 identified by LC-MS

| Protein name | MW(KDa) | Uniprot ID | Score in H4G-IP | Score in H4-IP |
| --- | --- | --- | --- | --- |
| Chromatin binding proteins |  |  |  |  |
| SP16H | 119.9 | Q9Y5B9 | 82 | 71 |
| SSRP1 | 81.1 | Q08945 | 52 | 69 |
| RBAP46 | 47.8 | Q16576 | 56 | 297 |
| RBAP48 | 47.7 | Q09028 | 56 | 451 |
| NASP | 85.2 | P49321 | 43 | 621 |
| ASF1A | 23 | Q9Y294 | N/A | 162 |
| CAF1B | 61.5 | Q13112 | N/A | 108 |
| CAF1A | 106.9 | Q13111 | N/A | 50 |
| Nucleolar proteins |  |  |  |  |
| NOP58 | 59.6 | Q9Y2X3 | 67 | N/A |
| NPM1 | 32.6 | P06748 | 70 | N/A |

**Figure 5.**
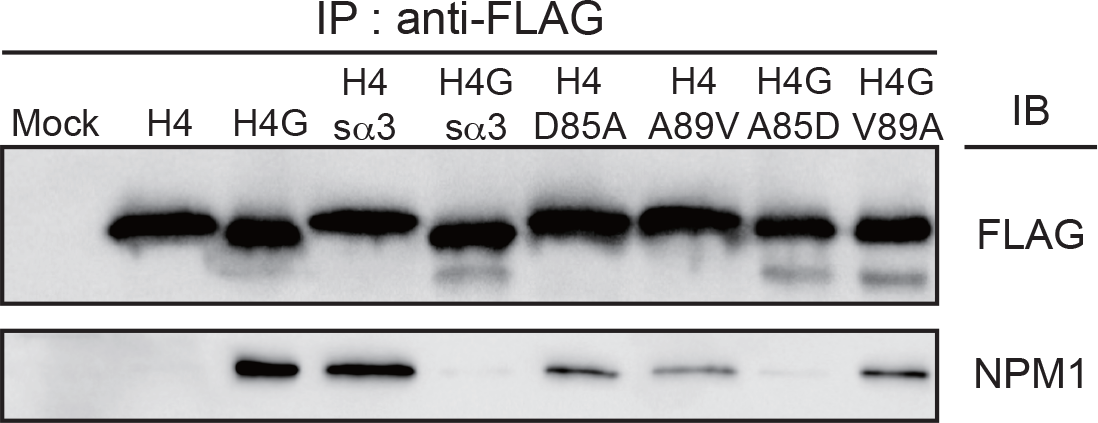
H4G strongly interacts with NPM1 and this interaction is mediated by μ-helix 3 domain. Immunoprecipitation using FLAG-proteins to identify the interaction of H4G with the nucleolar histone chaperone NPM1.

### H4G regulates the synthesis of rRNA and cell growth

We next assessed whether the nucleolar localization of H4G plays a role in rDNA transcription or ribosome biogenesis. To this end, we treated H4G-transfected MCF7 cells with a low concentration of actinomycin D, which primarily inhibits transcription by RNA polymerase I [34, 35], and observed that the nucleolar localization of H4G was lost with H4G dispersed throughout the nucleus (Figure EV5). In contrast, in non-transfected control cells, the addition of actinomycin D did not change the localization of H4. Similar results were obtained in other cell lines, including HeLa, LCC1, and LCC2 cells (data not shown). In addition, H4G localization did not change when H4G-transfected MCF7 cells were treated with rapamycin, an mTOR signaling inhibitor. mTOR signaling has been shown to regulate the synthesis of ribosome components, pre-rRNA, and 5S rRNA without affecting rDNA transcription [36] (Figure EV5). These results suggest that the nucleolar localization of H4G may be dependent on ongoing rDNA transcription.

Despite these findings, it remains unclear whether the nucleolar localization of H4G has a functional role in the regulation of rDNA transcription. To address this issue, and to study the functional role of H4G, we produced an H4G knockout in the MCF7 cell line using CRISPR-Cas9. We found that, in the H4G knockout cells, the amount of rRNA was reduced (Figure 6A). To examine whether transcription or RNA processing were affected by H4G levels, we compared rRNA levels by Q-PCR using primers for the external transcribed spacers (ETS) at the 5′ UTR of the rRNA. Our results obtained with the ETS primer were similar with that of the total rRNA (Figure 6A). To confirm H4G knockout is not caused by off-target effects, we produced the H4G rescue lines by transfecting the pLVX-TetOn-Puro H4G 3xFLAG plasmid into H4G knockout cells. The amount of ETS rRNA in the H4G rescue lines was similar to the amount observed in the WT MCF7 line (Figure EV6). These data indicate that H4G promotes the bulk transcription of rRNA.Because of the H4G nucleolar localization and its contribution to rRNA synthesis, we further hypothesized that H4G may affect protein synthesis in general. Therefore, we quantified the amounts of protein synthesis, as measured by OPP (O-propargyl-puromycin) incorporation, and found that protein synthesis rates were reduced by approximately 10% in H4G knockout cells compared with WT cells (*N* = 3; Figure 6B). Moreover, we quantified each cell cycle stage at 18 h after release from G_0_ (Figure 6C and 6D) and found that 30%–40% of the overall cell cycle delay was observed in the H4G knockout cells (Figure 6D). These results suggest that H4G is directly involved in rDNA transcription and affects cellular proliferation.

**Figure 6.**
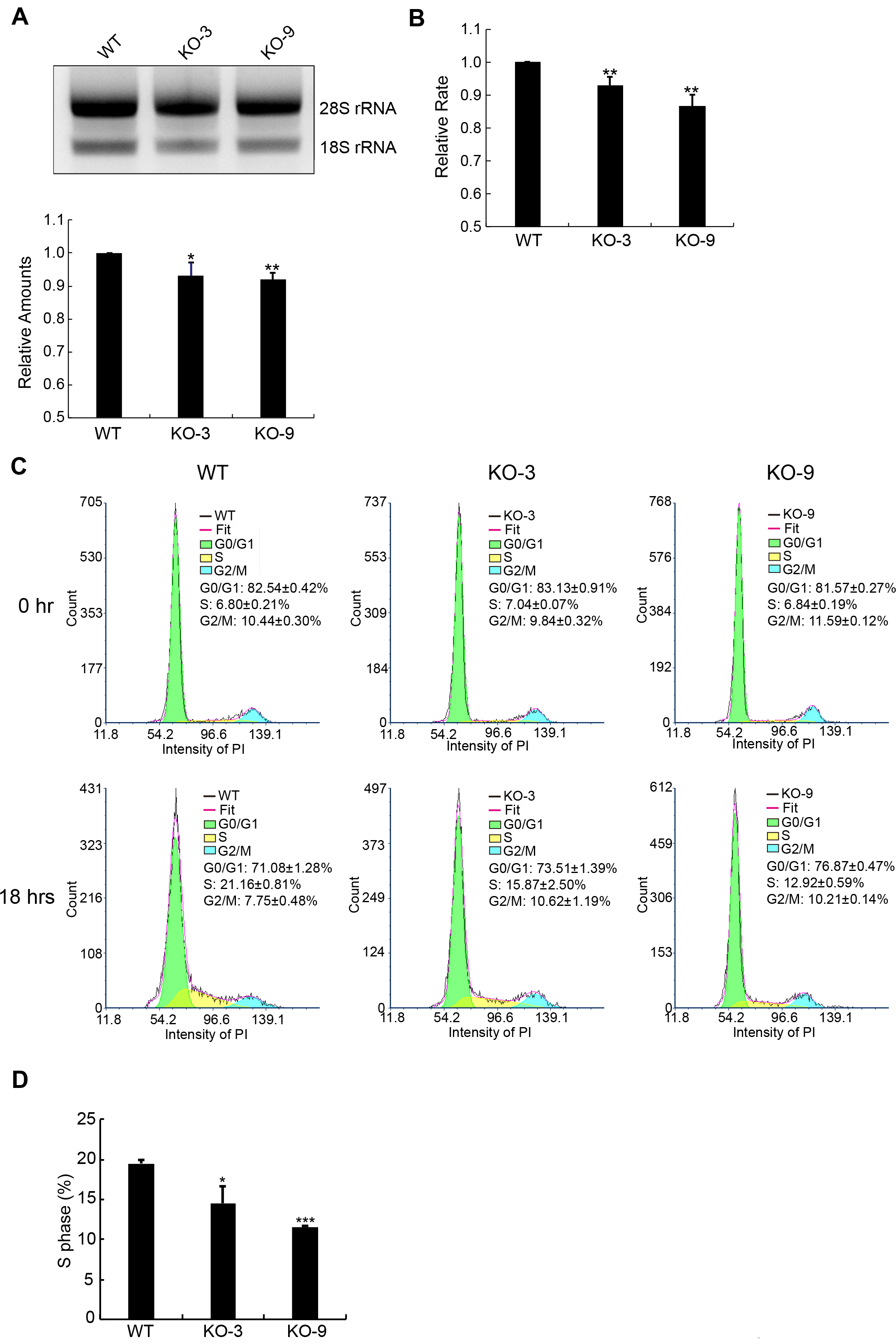
H4G is involved in rRNA transcription. (A) Comparison of rRNA amounts among WT and H4G knockout (H4GKO) MCF7 cell lines. Total RNA was normalized by GAPDH mRNA and loaded onto agarose gels. Error bars represent SD in triplicate experiments. *p < 0.05 and **p < 0.01 (Student’s *t*-test). (B) The relative protein synthesis rate among WT and H4GKO MCF7 cell lines. Error bars represent SD. **p < 0.01. (C) Cell cycle histograms acquired by flow cytometry and quantification of the subpopulation fraction of the histogram among WT and H4GKOs MCF7 cells. FACS plots and data are representative of at least three separate experiments. (D) The percentage of S phase cells among WT and H4GKOs MCF7 cells. Error bars represent SD. *p < 0.05, ***p < 0.001 (Student’s *t*-test).

## Discussion

In this study, we have characterized H4G, a novel hominidae-specific histone H4 variant. Although this variant has initially been reported to be a replication-dependent histone [18], we were able to obtain cDNA for H4G from polyA-mRNA. Indeed, the presence of poly A+ in histone mRNA is characteristic of replication–independent histone variants [17]. The cell cycle–independent, intron-containing, histone H4 gene *h4r* has been previously identified in *Drosophila*. However, it encodes a protein identical to that of the cell cycle–dependent H4 gene counterpart [37]. The reason(s) why *Drosophila* has cell cycle–dependent and cycle–independent H4 genes remains unclear. However, the situation is different in primate H4G, in which the amino acid sequence shares only 85% sequence identity with the canonical H4 counterpart. Furthermore, the *h4g* gene does not contain introns.

At the cellular level, H4G exhibits a nucleolar localization pattern and appears to interact with the nucleolar histone chaperone protein NPM1. Several other nucleolar histone chaperones have been identified, including factor Spt2 and nucleolin. Nucleolin directly binds to H2A-H2B dimers in order to facilitate nucleosome assembly [38], whereas the transcription factor Spt2 functions as a nucleolar histone chaperone by interacting with the H3/H4 tetramer [39, 40]. The Spt2 interaction with H4 involves the alpha 3 domain of H4 [39], which is a diverging sequence from that of H4G. Our H4G affinity interaction assay was not able to detect nucleolin nor Spt2, likely due to the stringent washing conditions used.

Of special interest is the expression of the *h4g* gene in cancer: we found *h4g* mRNA to be restricted to breast cancer cell lines and in breast cancer patient tissues. Our functional analyses further revealed that H4G is likely in rDNA transcription. One well-studied histone variant involved in cancer progression is H3.3 and its K27M mutation, which occurs in pediatric high-grade glioma [41, 42]. This mutation blocks the polycomb repressor complex 2 methylation activity of H3K27, which is a marker for gene silencing [43–46]. Indeed, the reduction of H3K27 trimethylation is associated with cancer-associated genes such as p16INK4A and CDK6 and contributes to tumorigenesis [47].

Other histone variants also show expression changes in human cancer progression, including macroH2A, H2A.Z, H2AX and H3.3 [for more details see the reviews by Vardabasso et al. [11] and Buschbeck and Hake [48]]. Indeed, the expression of H2A.Z is increased in colorectal, breast, lung, and bladder cancers. The elevated expression of H2A.Z is also significantly associated with metastasis to lymph nodes and shorter survival [11, 48]. However, for many of the histone variants, it is not clear whether altered expressions are a consequence of the cancer state or if they act as oncogenic factors. Even though H4G is a histone variant expressed in cancer patients, several differences are noteworthy. First, while the expressions of other histone variants are altered in cancer patients and cells, they are also present in normal cells whereas H4G appears to be preferentially expressed in breast cancer cell lines and breast tissues from breast cancer patients. Second, H4G is uniquely localized to the nucleoli, where it enhances rRNA expression. How *h4g* gene expression is triggered by tumorigenicity is intriguing and remains to be elucidated.

## Material and Methods

### Plasmids

Human H4, H4G, H4sN, H4GsN, H4GsC, H4GsN+C, H4sα1, H4sα2, H4sα3, H4Gsα3, H4D85A, and H4A85V sequences were inserted into the p3xFLAG-CMV-14 vector (Sigma-Aldrich). For stable cell line construction, the human H4G gene with the C-terminal 3xFLAG sequence was inserted into the pLVX-Tet-On-Puro vector (Clontech). For *in vitro* nucleosome reconstitution, the *h4* and *h4g* genes were inserted into the pGEX4T3 vector, and the canonical histones H2B, H2A, and H3 were cloned into the pET11a vector.

### *In vitro* nucleosome reconstitution and MNase digestion

The human histones H2A, H2B, H3, GST-H4 (gH4), and GST-H4G (gH4G) were expressed in *Escherichia coli* BL21-codonplus (DE3), and they were purified from the inclusion bodies [49]. The H3-gH4, H3-gH4G, and H2A-H2B complexes were reconstituted as follows: Purified gH4 or gH4G was mixed with H3 at a 1:1 molar ratio in unfolding buffer (20 mM Tris–HCl [pH 7.5], 7 M guanidine hydrochloride, and 20 mM 2-mercaptoethanol), and the mixture was dialyzed against refolding buffer (10 mM Tris–HCl [pH 7.5], 2 M NaCl, 1 mM EDTA, and 5 mM 2-mercaptoethanol). The H3-gH4/H3-gH4G and H2A-H2B complexes were mixed at a molar ratio of 4:1 and further mixed with the 206bp DNA containing the 601 nucleosome positioning sequence in a solution containing 2 M NaCl. The nucleosomes were then reconstituted by the salt-dialysis method [30, 50], and 20 μL of the reconstituted nucleosome was digested by the indicated amount of MNase (Worthington) at room temperature for 5 min.

### Cell culture

The MCF-7, LCC1, LCC2, MCF-10A, and HEK293T cell lines were purchased from ATCC and cultured in Dulbecco’s modified Eagle’s medium supplemented with 10% fetal bovine serum and 1% penicillin/streptomycin at 37°C with 5% CO_2_ . Plasmid transfection was performed using Lipofectamine 2000 (Thermo Fisher Scientific) or Polyethylenimine (Polysciences).

### Immunoprecipitation

HEK293T cells transfected with the according plasmids were collected 48 h after transfection. For immunoprecipitation, cell extracts were prepared at 4°C in a lysis buffer containing 10 mM HEPES pH 7.9, 1.5 mM MgCl_2_, 10 mM KCl, 420 mM NaCl, 0.5% NP-40, 5 mM 2-mercaptoethanol, and protease inhibitor cocktail (Roche) with sonication. Anti-FLAG M2-coupled beads (Sigma-Aldrich) were added to the extracts and incubated at 4°C for 2 h. Subsequently, the beads were extensively washed in lysis buffer, and the bound proteins were analyzed by immunoblotting against FLAG (FLAG M2 antibody from Sigma-Aldrich) and NPM1 (Clone FC82291 from Abcam).

### Mass spectrometry analysis and data processing

Anti-FLAG immunoprecipitation was performed as described above. After immunoprecipitation, bound proteins were eluted by incubating the beads with the FLAG peptide (1 μg μL^−1^). Eluted proteins were resolved using SDS-PAGE and stained with SYPRO Ruby (Thermo Fisher Scientific). For protein identification, gels that contained the interacting proteins were cut and analyzed using an LTQ Velos liner ion-trap LC-MS system in HKUST BioCRF. The acquired tandem mass spectra were then subjected to gene database searches using the MASCOT search engine (Matrix Science).

### Knockout and rescue cell line establishment

To construct a knockout H4G line in MCF7 cells, sgRNA (5′-CACCGTTCGGGGCAAGGCCGGAAA-3′) that targets HIST1H4G was inserted into the PX459 plasmid (Addgene). Transfection plasmids in MCF7 cells were carried out with the FuGENE^®^ HD Transfection Reagent (Promega) according to the manufacturer’s instructions. Genomic DNA was extracted and subjected to PCR using a forward primer (5′-GGACGAATTCTCCCGCCTTTCCTGGTCTTTCAG-3′) and reverse primer (5′-GTTAGGATCCCAGGGTTCTTCCCTGGCGTT-3′) to generate a product spanning the targeted region. The pLVX-TetOn-Puro H4G 3xFLAG plasmid was then transfected into the KO cell lines and selected with puromycin for rescue cell line establishment.

### RNA extraction and quantitative real-time PCR

RNA isolation was performed using the PureLink RNA Mini Kit (Invitrogen). cDNA was generated using Superscript III First-Strand Synthesis System (Thermo Fisher Scientifics) using a random hexamer primer. A negative control was generated by replacing reverse transcriptase with water in the cDNA synthesis process to exclude genomic DNA contamination. Messenger RNA (mRNA) expression was measured using quantitative real-time PCR (q-RT-PCR) assays using gene-specific primers (SYBR Green assay) by a LightCycler^®^ detector (Roche). The relative fold change for each gene was calculated using the ΔΔCt method as previously described **[51]** and the Student’s *t*-test was used to determine statistical significance. Primers used for q-RT-PCR were H4G forward (5′-TTTAGAGATAGTTCTGACTTGT-3′) and reverse (5′-AGGAAAGGCCTGGCCTCACTTA-3′); GAPDH forward (5′-CTCCTGCACCACCAACTGCT-3′) and reverse (5′-GGGCCATCCACAGTCTTCTG-3′); and ETS forward (5′-GAACGGTGGTGTGTCGTT-3′) and reverse (5′-GCGTCTCGTCTCGTCTCACT-3′). For quantification of rRNA, total RNA was normalized by *GAPDH* mRNA and loaded on agarose gels. The 18S and 28S rRNA were quantified using ImageJ software.

### Protein synthesis assay

MCF7 cells were stained with the Click-iT^®^ Plus OPP Alexa Fluor 488 Protein Synthesis Assay Kit (Life Technologies). OPP was added at a 10 μM final concentration, and cells were incubated for 30 min. Cells were then fixed with 70% ethanol and processed as instructed in the OPP assay kit. Cells were analyzed using flow cytometry.

### Cell cycle analysis

MCF7 cells were synchronized by serum starvation for 24 h and induced to re-enter the cell cycle by the addition of serum. Cells were then harvested for propidium iodide staining and analyzed by fluorescence-activated cell sorting (FACS) using a Becton Dickinson FACSAria™ III flow cytometer (BD Biosciences) to determine the cell cycle fraction. Data were analyzed using FCS Express software (De Novo Software).

## Acknowledgements

This work was supported by grants from the Research Grants Council of the Hong Kong SAR (16104917, 26100214, C-702915G) to TI and by Canadian Institutes of Health Research (CIHR) grant (MOP-130417) to JA.

## Conflicts of interest

The authors declare no conflicts of interest with the contents of this article.

## Author contributions

M.L., X.S., J.A., Y.Y., and T.I. designed the experiments. M.L., X.S., W.S., and Y.A. performed the experiments. T.L., D.D., and M.S.C provided the material for the experiments. N.M. and C.N. provided information necessary for the publication. M.L., X.S., J.A., Y.Y., and T.I. wrote the manuscript.

**Figure EV1.** The comparison of hydrophobicity between human H4 and H4G using Expasy ProtScale with the Kyte & Doolittle method.

**Figure EV2.** The sub-nucleolar localization of H4G and H4GsN with the (**A**) fibrillanin antibody and the (**B**) nucleolin antibody.

**Figure EV3.** Immunoprecipitation using FLAG-proteins H4, H4G, and H3 to quantify the interaction with NPM1.

**Figure EV4.** Representative SDS-PAGE gel of mass spectrometry samples for detecting H4-and H4G-interacting proteins. Asterisk (*) represents either the H4-FLAG or the H4G-FLAG protein.

**Figure EV5. Immunostaining of MCF7 cells with the addition of actinomycin D (400 nM) or rapamycin (100 nM) for 4 h.**

**Figure EV6. Quantification of relative rRNA amounts among WT, H4GKO, and H4G rescue MCF7 cell lines using the external transcribed spacers (ETS) region at the 5′ UTR primer.** Data were normalized to GAPDH. Error bars represent SD in triplicate experiments. *p < 0.05 and **p < 0.01 (Student’s *t*-test).

